# High-throughput genotyping of the spelt gene pool reveals patterns of agricultural history in Europe

**DOI:** 10.1101/481424

**Authors:** Michael Abrouk, Christoph Stritt, Thomas Müller, Beat Keller, Anne C. Roulin, Simon G. Krattinger

**Author notes:** contributed equally (Michael Abrouk, Christoph Stritt, Thomas Müller).

## Abstract

Spelt, a close relative of hexaploid bread wheat and a dominant wheat subspecies cultivated in Europe before the 20^th^ century, still plays an important role as a high-value niche product today. Compared to most other cereals, spelt has not been subjected to intensive breeding in the 20^th^ century. Even today, mostly traditional landraces are cultivated on a regional scale. The traditional way of spelt cultivation has limited the extensive exchange of germ plasm and intermixing of genetic material, which makes spelt an ideal crop to study the early agricultural history of cereals in Europe. Here, we unraveled the population structure and agricultural history of spelt based on 22,999 high-quality SNPs obtained by genotyping-by-sequencing on 267 spelt accessions covering the entire cultivation range. SNP markers were aligned to the high-quality bread wheat reference genome, which allowed us to analyze individual subgenomes. Our analyses of genetic variation revealed that bread wheat and spelt are most likely of monophyletic origin, but that European spelt diverged from bread wheat by hybridization with tetraploid emmer wheats. Interestingly, spelt accessions from the Iberian Peninsula formed a separate clade that was distinct from the Central European accessions for all three subgenomes. Demographic modelling indicated that Iberian spelt was introduced into Europe independently from Central European spelt. Our analysis provides a comprehensive assessment of spelt diversity and history. The separate introduction of Iberian spelt is supported by recent molecular evidence of two independent prehistoric migrations of ancient farmers from the Near East into Europe.

## Introduction

Recent genomic advancements have allowed to shed light on the population structures and domestication histories of different crop species (Cubry et al., 2018; W. S. Wang et al., 2018; G. A. Wu et al., 2018; J. Wu et al., 2018). The domestication of wild ancestors of today’s crops started around 10,000 years ago and resulted in the gradual adaptation of wild plants to agricultural practices. Plant domestication was not only associated with profound sociocultural changes during the Neolithic Revolution, e.g. the transition from hunter-gatherers to sedentary communities, but also led to distinct morphological alterations in domesticated plants compared to their wild relatives. The ‘domestication syndrome’ includes loss of natural seed dispersal mechanisms (non-shattering), increase of fruit or seed size, and changes in plant architecture (Meyer, DuVal, & Jensen, 2012). The adaptation of domesticated plants to agricultural practices and human needs was further accelerated after the description of the laws of genetic inheritance in the 19^th^ century, which laid the foundation for modern plant breeding of the 20^th^ century. As a consequence, today’s elite crop cultivars greatly outperform their wild ancestors with regards to yield, quality, and adaptation to a wide range of climatic conditions. On the other hand, domestication, early landrace selection, and modern breeding in particular have reduced the genetic diversity in the gene pools of many crops (Tanksley & McCouch, 1997; Wulff & Dhugga, 2018). The introduction of genetically heterogeneous landraces into modern breeding programs is a valuable approach to counteract erosion of genetic diversity (Dwivedi et al., 2016). The hundreds of thousands of crop accessions maintained in germplasm collections worldwide thus play an important role in providing solutions to some of the most pressing challenges faced by agriculture, including climate change and population growth (Wambugu, Ndjiondjop, & Henry, 2018).

Crop domestication occurred independently at different locations. One of the domestication hotspots is the Fertile Crescent, a region in the Near East that spans today’s Jordan, Israel, Lebanon, Syria, Iraq, southeast Turkey, and western Iran (Salamini, Ozkan, Brandolini, Schafer-Pregl, & Martin, 2002). The Fertile Crescent is the center of origin of some of the most important crop plants, including wheat, barley, rye, lentil, chickpea, and pea. Following domestication, these crops were introduced into different parts of the world, most likely along with human migration. Wheat is one of the most important cereal crops that serves as a staple food for more than two billion people (International Wheat Genome Sequencing Consortium, 2014). The two most widely cultivated wheat species today are tetraploid durum wheat, also called pasta wheat (*Triticum durum*), and hexaploid bread wheat (*T. aestivum* ssp. *aestivum*). A close relative of bread wheat is hexaploid spelt (*T. aestivum* ssp. *spelta*) that is mainly cultivated in Central Europe and northern Spain as a high-value niche product. Spelt, however, was a staple crop in Europe from the Bronze Age until about the beginning of the 20^th^ century (Muller et al., 2018). In some regions of Central Europe, particularly in southern Germany and Switzerland, spelt even replaced tetraploid emmer as the principal wheat species in the early Iron Age (around 750 BC) (Nesbitt & Samuel, 1996). In 1930, spelt still accounted for about 40% of the Central European wheat production area (Kema, 1992). Despite being a domesticated wheat, spelt shows some morphological characteristics that resemble non-domesticated grass species, including a brittle rachis and kernels that are tightly surrounded by tenacious glumes. Because of the brittle rachis, spelt spikes disintegrate into individual spikelets more easily than bread wheat spikes (shattering) and the tenacious glumes require vigorous processing to free the grains (Figure 1). While these characteristics are beneficial for the dispersal and protection of seeds in wild plants, they are undesirable for mechanical harvesting and processing, which is one of the main reason for the replacement of spelt with free-threshing bread wheat in the 20^th^ century. The two hexaploid wheat subspecies *aestivum* and *spelta* can be freely intercrossed, which breeders continue to exploit to transfer agronomically important genes from spelt into the bread wheat gene pool (Dyck & Sykes, 1994; Sun, Wei, Ni, Xie, & Yang, 2002).

**Figure 1.**
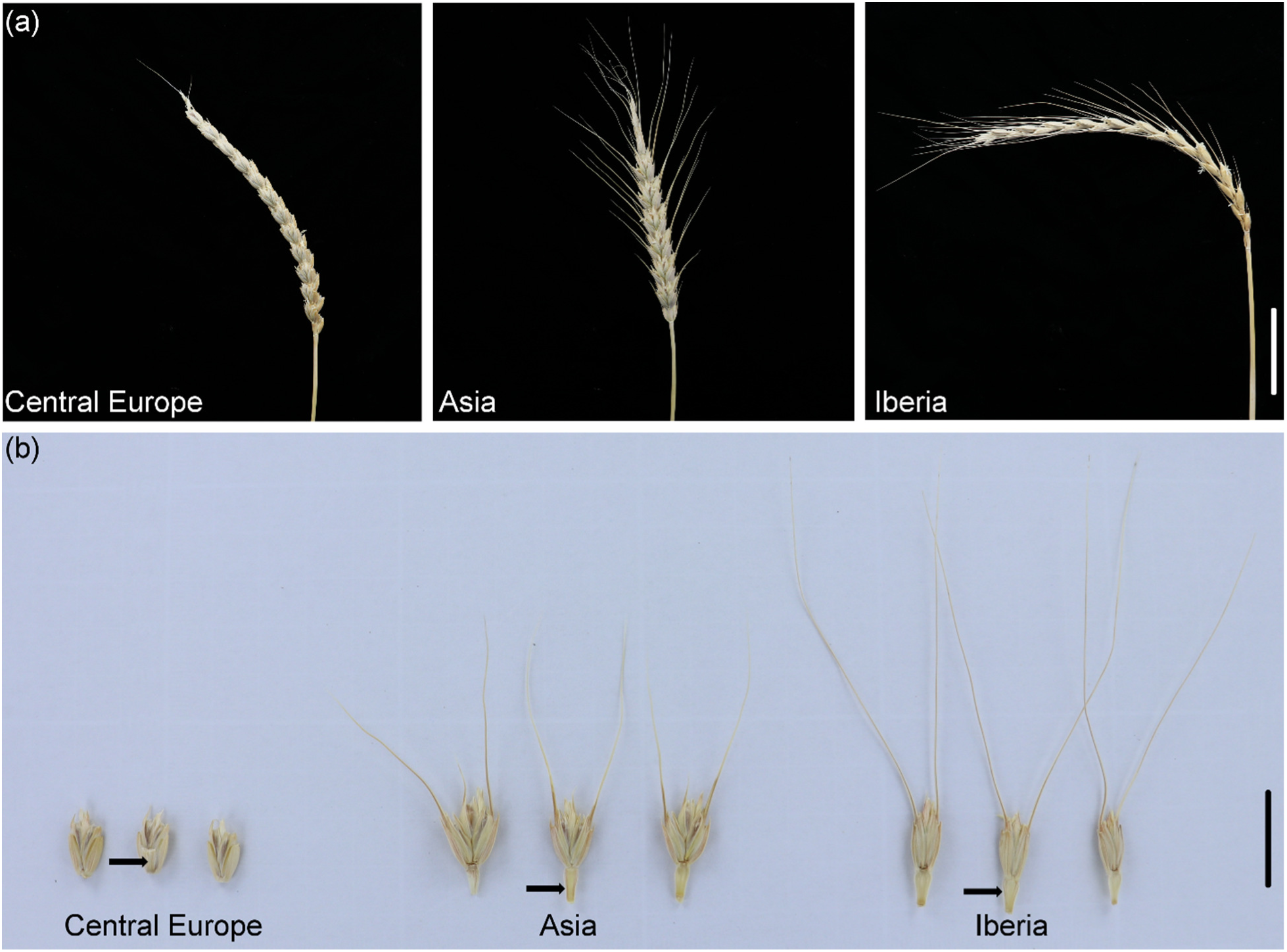
Spike morphology and spikelet disarticulation in spelt. (a) Representative spikes of spelt accessions collected in Central Europe, Asia, and the Iberian Peninsula. Scale bar = 5 cm. (b) Spikelet disarticulation of spelt. Spikelet disarticulation in plants with a brittle rachis can be classified as barrel-type (upper rachis segment pressed against the lower spikelet) and wedge-type (rachis segment pointing down). Central European spelt shows a barrel-type disarticulation, whereas disarticulation in Asian and Iberian spelt is of the wedge type. Scale bar = 2 cm. Arrows point to the rachis segment after spikelet disarticulation.

Hexaploid wheat emerged after two independent hybridization events that involved three diploid wild grass species of the Triticeae tribe. The first hybridization between wild einkorn (*T. urartu*, AA genome) and a close relative of *Aegilops speltoides* (BB genome) formed tetraploid wild emmer (*T. turgidum* ssp. *dicoccoides*, AABB genomes). Bread wheat arose after a second hybridization of domesticated emmer (*T. turgidum* ssp. *dicoccum*, AABB genomes) with the D genome of wild goatgrass (*Ae. tauschii*). The first hybridization occurred a few hundred thousand years ago, while the second hybridization is thought to have occurred in a field of domesticated emmer around 10,000 years ago (Marcussen et al., 2014). While both wild and domesticated forms of diploid and tetraploid wheats are described, no wild hexaploid wheat exists. Despite the close relatedness of spelt and bread wheat, the population structure and agricultural history of spelt is still ambiguous and has not been well studied on a whole-genome level. Phylogenetic analyses based on a few microsatellite markers or single gene sequences suggested three spelt clades – an Asiatic, a Central European and an Iberian clade (Bertin, Gregoire, Massart, & de Froidmont, 2004; Blatter, Jacomet, & Schlumbaum, 2004; Dvorak et al., 2012; Elia, Moralejo, Rodriguez-Quijano, & Molina-Cano, 2004). Two observations made by these studies are of particular interest. Comparisons of the A and B subgenomes indicated a polyphyletic origin of European spelt and bread wheat, while comparisons of the D subgenome suggested a common, monophyletic origin. Asiatic spelt is closer to bread wheat than to European spelt. Based on these findings, it was suggested that bread wheat and spelt have a common origin in the Fertile Crescent. During its migration to Europe, a free-threshing hexaploid wheat might have hybridized with a hulled tetraploid emmer wheat, which gave rise to European spelt (Blatter, Jacomet, & Schlumbaum, 2002; Dvorak et al., 2012; Liu & Tsunewaki, 1991). This could explain why the A and B subgenomes of European spelt markedly differed from bread wheat, while their D subgenomes are more similar. Tetraploid and hexaploid wheats were often cultivated in the same fields, which would have facilitated spontaneous hybridizations. However, these studies probed the genome with a very low number of non-randomly distributed molecular markers, which did not allow to get a comprehensive, genome-wide insight into the population structure of spelt. With almost 16 billion base pairs (16 Gb) the genome of hexaploid wheat is exceptionally large (International Wheat Genome Sequencing Consortium, 2018). Whole-genome re-sequencing of many wheat accessions for population genomic analyses is thus not yet a feasible strategy and genotyping-by-sequencing (GBS) is frequently used to comprehensively genotype large wheat collections (Poland, Brown, Sorrells, & Jannink, 2012; Schreiber, Stein, & Mascher, 2018). The recent completion of a high-quality wheat reference sequence now allows to anchor GBS markers and to investigate individual subgenomes and specific genomic regions (International Wheat Genome Sequencing Consortium, 2018).

Here, we report on the genotyping of 267 spelt accessions collected from the entire range of spelt cultivation. This comprehensive analysis of the spelt gene pool allowed us to unravel population structures and agricultural history with genome-wide marker coverage.

## Materials and Methods

### Plant material

Seeds of 75 *T. aestivum* ssp. *aestivum* accessions and 267 *T. aestivum* ssp. *spelta* accessions used in this study were obtained from four germplasm collections (the national genebank of Agroscope, Switzerland; the genebank of IPK Gatersleben, Germany; the Centre for Genetic Resources, Wageningen University, the Netherlands; USDA National Plant Germplasm System, USA). The spelt accessions where chosen to cover the entire cultivation range of the subspecies. Twenty seven accessions derive from crosses between bread wheat and spelt in recent breeding programs. The set includes spelt from Central Europe, Southern Europe, Africa, Asia, and America. The *T. aestivum* ssp. *aestivum* accessions all originate from Central Europe (Supporting Information Table. S1).

### DNA extraction

Single plants were grown and leaf segments of approximately 3 cm were sampled from the first leaf of 12-day-old seedlings. Frozen leaf tissue was grinded with a Mixer Mill MM 400 (Retsch, Haan, Germany) for 60 sec at 13,000 rpm. Grinded leaf tissue was suspended in 500 μl extraction buffer (1 M guanidine thiocyanate, 2 M sodium chloride, 30 mM NaAc, pH 6.0, 0.2 % Tween 20) and incubated at 65 °C for 30 min. The lysate was centrifuged for 10 min at 2,500 rpm and 300 μl of the supernatant was transferred onto DNA spin columns (Epoch Life Science, Inc, Texas, US). The columns were centrifuged at 13,000 rpm for 1 min to bind the DNA, washed twice with 600 μl and 250 μl wash buffer (50 mM sodium chloride, 10 mM Tris/HCl, pH 8.0, 1 mM EDTA in 70% ethanol) and centrifuged again at 13,000 rpm for 1 min after each wash step. The DNA was eluted from the silica columns with 100 μl TE buffer (10 mM Tris/HCl, pH 8.0, 1 mM EDTA) into new tubes.

### Genotyping by sequencing

DNA samples were genotyped with genotyping-by-sequencing (GBS) using *PstI* as restriction enzyme by the Institute of Biotechnology, Cornell University, NY, USA. The samples were sequenced on six lanes of a HiSeq 2000 and six lanes of a NextSeq 500 yielding 1,915,955,758 reads in total.

### SNP calling

Reads were mapped against the reference genome of Chinese Spring (IWGSC RefSeq v1.0) (International Wheat Genome Sequencing Consortium, 2018) with bwa v0.7.15 (Li & Durbin, 2009) and SNPs were called using the TASSEL v5.2.31 GBS pipeline (Glaubitz et al., 2014) using a kmer length of 64bp. SNPs were filtered using vcftools v0.1.14 (Danecek et al., 2011) based on the following criteria: (1) variant should be a biallelic SNPs, (2) SNPs with more than 20% missing data were discarded, (3) genotypes having more than 20% missing information were discarded and (4) SNPs with MAF ≥ 0.05 were retained (Eltaher et al., 2018). Our final set of SNPs consisted of 23,554 SNPs represented in 339 accessions.

### Principal Component Analysis

Principal Component Analyses (PCA) were performed with a homemade script in Python v2.7.12 using the libraries scipy v0.19.1, NumPy v1.13.3 (van der Walt, Colbert, & Varoquaux, 2011), sklearn v0.17.1 (Pedregosa et al., 2011), pandas v0.20.3 (McKinney, 2010), matplotlib v2.1.0 (Hunter, 2007) and ipython v6.1 (Perez & Granger, 2007).

### SNPhylo

We generated a maximum likelihood (ML) tree using the SNPhylo software package (Lee, Guo, Wang, Kim, & Paterson, 2014) with the 22,999 anchored SNPs. Low quality SNPs based on allele frequency (MAF ≤ 0.05), missing data (≥ 20%) and linkage disequilibrium (LD ≤ 1) were removed using the SNPRelate package (Zheng et al., 2012). Multiple sequence alignment of the concatenated SNPs for each sample was done using MUSCLE (Edgar, 2004) and a phylogenetic tree was constructed by running DNAML from the PHYLIP package (Felsenstein, 2005). One hundred bootstraps using the phangorn package (Schliep, 2011) were done and the tree was visualized and annotated with FigTree v1.4.3 program.

### Genetic clustering analysis

In order to estimate genetic clustering among the accessions, a structure analysis was performed with the ADMIXTURE software (Alexander, Novembre, & Lange, 2009). This type of analysis infers the population structure for a number of ancestral populations (K), as defined by allele frequencies at all studied loci, and simultaneously assigning ancestry proportions (or probabilities) to each individual in the study. We used the dataset of 22,999 SNPs from the 339 accessions. Data management and quality control operations were performed using vcftools v0.1.14 (Danecek et al., 2011) and PLINK v1.9 (Purcell et al., 2007). We explored K values from 2 to 10 and determined the lowest cross-validated error rate.

### Demographic inference

Demographic models were fitted to folded 2D site frequency spectra with fastSimcoal2 (version 2.6.0.3) (Excoffier, Dupanloup, Huerta-Sanchez, Sousa, & Foll, 2013). Accessions displaying discrepancies between their indicated geographic origin and the PCA cluster were removed. Only SNPs on the D subgenome were considered because of the suspected polyphyletic origin of the A and B subgenomes. We used biallelic SNPs filtered for missing data, as described above, but only removed singletons instead of sites with a MAF < 0.05. Since fastSimcoal assumes sites to be neutral and unlinked, we removed genic SNPs and only retained SNPs at least 20 kb apart from each other, leaving a total of 2,644 SNPs distributed among 209 spelt accessions from Asia, Central Europe, and Iberia.

Folded two-dimensional site frequency spectra were estimated with dadi (Gutenkunst, Hernandez, Williamson, & Bustamante, 2009). The numbers of alleles were projected down from 36 to 24, from 324 to 254, and from 58 to 42 for the Asian, the Central European, and Iberian spelt respectively, so as to maximize the number of segregating sites used to construct the SFS. A mutation rate of 1.3×10^−8^ (Ma & Bennetzen, 2004) was assumed. Further model specifications and parameter search ranges can be found in Supporting Information Table S2 and Online Resource 1. For each of the six models, 20 independent runs with 50 ECM cycles and 100,000 simulations per estimation step were performed. Confidence intervals were obtained with a parametric bootstrap approach in which 100 data sets were simulated with the parameter values of the best model, and parameter values then inferred from the simulated data.

## Results

Most of the European spelt accessions analyzed in this study were collected by expeditions during the 1930s. While European spelt was well documented at the beginning of the 20^th^ century, the existence of spelt in Asia was only discovered in the late 1950s (Kuckuck & Schiemann, 1957; Luo, Yang, & Dvorak, 2000). Interestingly, many spelt accessions collected from the Iberian Peninsula were morphologically more similar to Asian spelt than to Central European accessions (Figure 1). For example, spikes of many Spanish and Asian spelt accession carry awns (needle-like structures formed at the end of the floret) (Caballero, Martin, & Alvarez, 2007), while Central European spelt is generally awnless (Figure 1a). Also, the spikelet disarticulation, which refers to the breakpoint of the rachis in relation to the spikelet, differed between spelt from the Iberian Peninsula and Central Europe (Figure 1b). Awn formation and spikelet disarticulation in wheat is controlled by only a few loci (Katkout et al., 2014; Yoshioka et al., 2017). The apparent morphological difference between Central European spelt and Iberian spelt does thus not necessarily reflect profound genomic variation. We therefore used GBS to assess the genetic variation of 267 spelt accessions. For a comparison, we also included 75 bread wheat accessions, most of them representing landraces from Europe (Supporting Information Table S1). In total, the GBS returned 255,396 SNPs that were anchored to the reference sequence of the bread wheat landrace Chinese Spring (International Wheat Genome Sequencing Consortium, 2018). Filtering for bi-allelic SNPs, missing data of ≤20%, and minor allele frequency (MAF) ≥5% resulted in the final set of 22,999 high-quality SNPs with known chromosome location, 8,878, 10,528, and 3,593 for the A, B, and D subgenomes, respectively (Table 1). Three spelt accessions were removed during the filtering because they had too many missing SNPs. The SNP markers were distributed across the entire genome, but were preferably located in gene-rich, telomeric regions (Supporting Information Figure S1). Of the 22,999 SNPs, 2,434 were located in coding sequences and 20,565 in intergenic regions or introns.

**Table 1.** Distribution of filtered SNP markers per chromosome. Unknown means that the SNPs were assigned to scaffolds in the Chinese Spring reference sequences that were not anchored to a chromosome.

|  | A genome | B genome | D genome | unknown |
| --- | --- | --- | --- | --- |
| **total** | **8,878** | **10,528** | **3,593** | **555** |
| chr 1 | 1,092 | 1,415 | 448 |  |
| chr 2 | 1,206 | 1,672 | 703 |  |
| chr 3 | 1,311 | 1,795 | 540 |  |
| chr 4 | 1,302 | 1,076 | 261 |  |
| chr 5 | 1,345 | 1,489 | 551 |  |
| chr 6 | 1,041 | 1,537 | 478 |  |
| chr 7 | 1,581 | 1,544 | 612 |  |

### Whole genome analysis

Principal component analysis (PCA), phylogenetic analysis, and genetic clustering algorithms consistently revealed a separation of spelt accessions into three distinct gene pools comprising accessions collected in Asia, Central Europe, and the Iberian Peninsula (Figure 2). Spelt accessions originating from Asia clustered together with the bread wheat accessions, while spelt accessions from Central Europe and the Iberian Peninsula formed two distinct groups. In the PCA, the first principal component mainly separated European spelt from bread wheat and Asian spelt. Although the bread wheat accessions used in this study originated from Central Europe, they were closer to Asian spelt than to the Central European spelt accessions. The second axis separated Iberian spelt from the other two gene pools. Together, the first two principal components explained 17.93% of the total variation (Figure 2a). Accessions that originated from artificial crosses between Central European spelt and bread wheat located in the middle of the first axis, confirming the accuracy of the sequencing and SNP calling. Two spelt accessions collected in America also localized within the wheat-spelt crosses in the PCA, indicating a hybrid origin. On the other hand, four accessions collected in Africa (one from Morocco and three from Ethiopia) clustered with the Central European spelt, possibly indicating that they were brought into Africa from Europe. Twenty-three spelt (8.7%) and one bread wheat (1,3%) accession showed discrepancies between their indicated geographic origin and the PCA cluster (Supporting Information Table S1). The most likely reasons for this are erroneous passport information or mistakes during propagation of genebank material. Alternatively, this pattern might reflect interchange of germ plasm between different regions before collection. A maximum-likelihood tree confirmed the PCA result and clearly visualized the three different spelt gene pools (Figure 2b). Inference of the population structure was done using the ADMIXTURE software assuming various ancestral populations K, ranging from 2 to 10. The split into the three main gene pools revealed by PCA and phylogenetic analyses was apparent at K=3 (Figure 2c). The minimal cross-validation error value was obtained at K=7 (CV = 0.57030; Supporting Information Figure S2), which indicates the most probable number of ancestral populations. Even with increasing K, the Iberian spelt remained a unique, homogenous cluster that did not split further. Compared to K=3, the Central European spelt accessions were further divided into four different groups at K=7, reflecting spring spelt, northern and southern accessions from Germany, and Switzerland. At K=6, a split of Asian spelt into western (Azerbaijan and Iran) and eastern accessions (Iran, Afghanistan and Tajikistan) became apparent. Despite the clear morphological differences in terms of spike shape, threshability and seed shattering between Asian spelt and bread wheat, the molecular analyses do not necessarily justify a separation into two subspecies.

**Figure 2.**
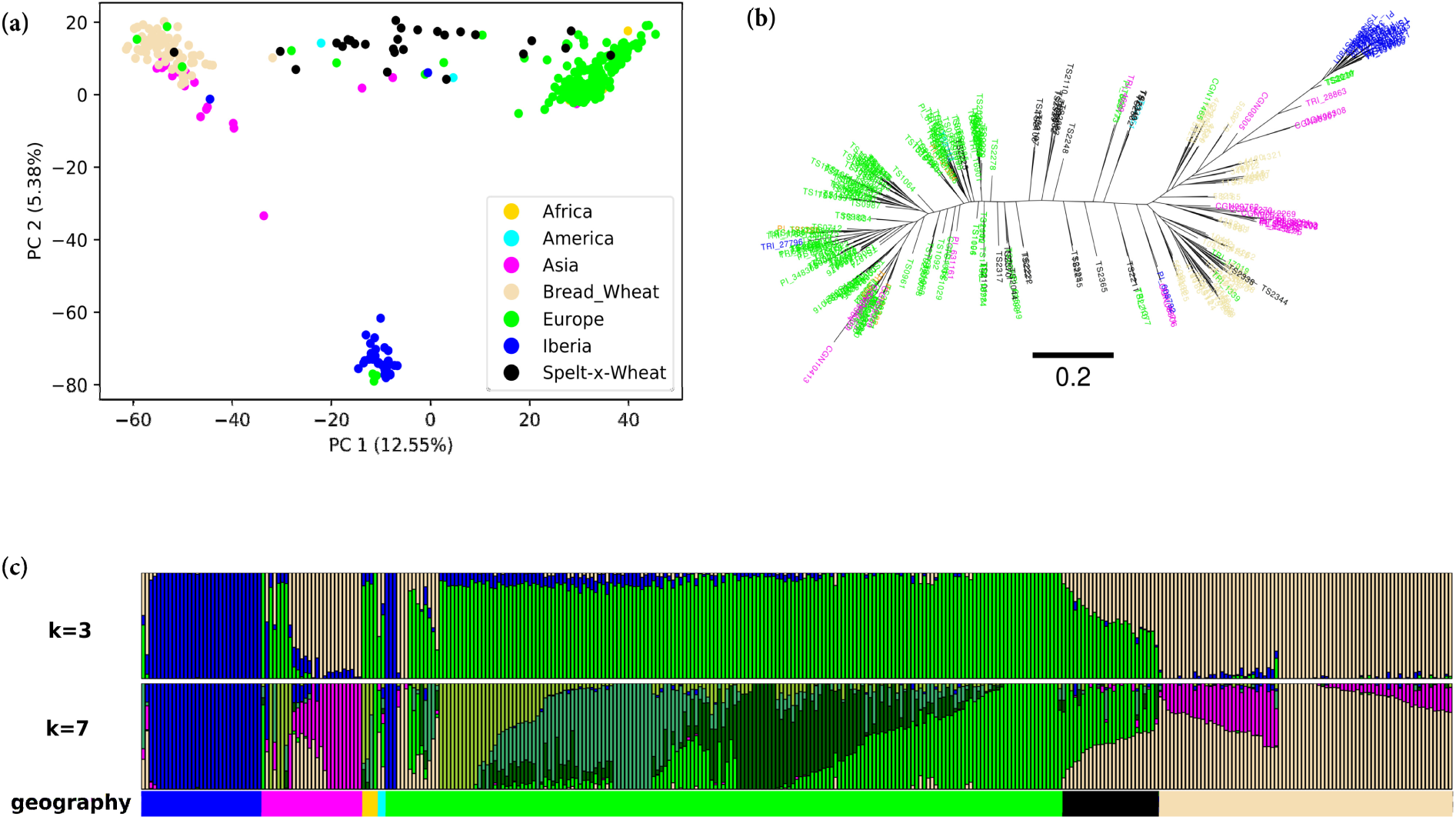
Whole genome analysis of spelt and bread wheat. (a) Principal component analysis (PCA) across the 339 spelt and bread wheat accessions based on 22,999 high-quality SNPs. Samples are colored according to subspecies and geographical origin. (b) Maximum-likelihood tree constructed with SNPhylo. The colors used to label the accessions are identical to (a) and the branch size is indicated below the tree. (c) ADMIXTURE ancestry coefficients (K=3 and K=7) for the 339 accessions of spelt and bread wheat. Stacked bars represent accessions and colors represent ancestry components. Accessions are ordered according to subspecies and geographical origin.

### Subgenome analysis

The same analyses as described above were performed individually for each of the three subgenomes, with 8,878 A, 10,528 B and 3,593 D subgenome SNPs, respectively (Figure 3).

**Figure 3.**
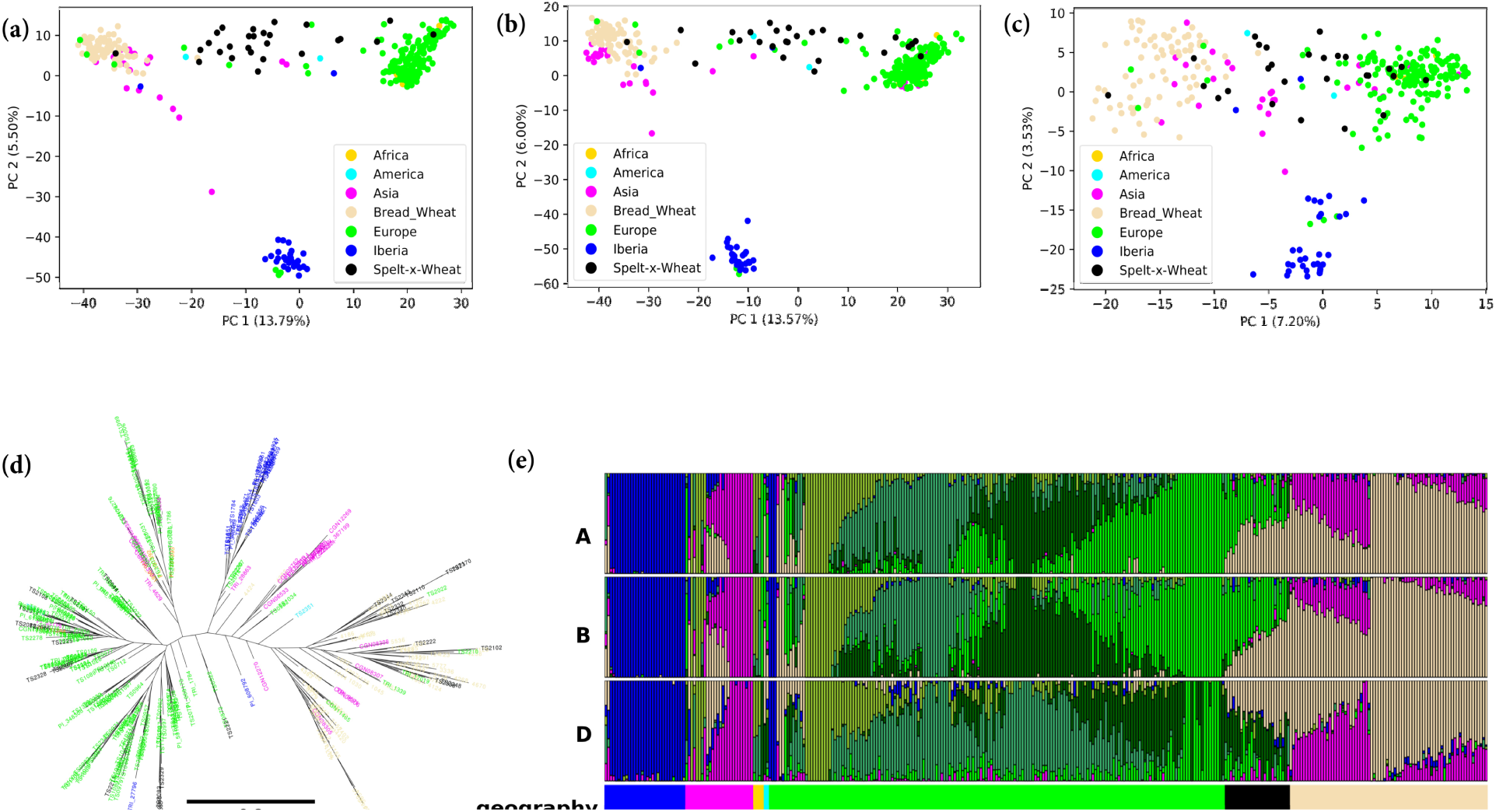
Subgenome analysis of spelt and bread wheat accessions. Principal component analyses for the A subgenome (a), B subgenome (b) and D subgenome (c) are shown. (d) Maximum-likelihood tree of the subgenome D. In both, PCA and phylogenetic tree, the colors represent the species and their geographical origin for spelt and the branch size is indicated below the tree. (e) ADMIXTURE ancestry coefficients (K=7) for the three subgenomes. Stacked bars represent accessions and colors represent ancestry components. Accessions are arranged according to the species and their geographical origin.

PCA, phylogenetic tree, and ADMIXTURE analyses of the A and B subgenomes revealed a picture very similar to the whole-genome analysis (Figure 3a,b; Supporting Information Figure S3) with a clear separation into a Central European, Asian, and Iberian gene pool. The situation was slightly different for the D subgenome (Figure 3c,d). While a separation into Central European, Iberian, and Asian spelt was still observed in the PCA, the distribution of the genotypes was more compressed. The total variation explained by the two first principal components (10.73%) was lower for the D subgenome compared to the A (19.29%) and B (19.57%) subgenomes, respectively. Similarly, the distances between the three clusters in the D subgenome maximum-likelihood tree were shorter compared to the A and B subgenomes, indicating a lower number of substitutions per site (Figure 3d; Supporting Information Figure S3). These results indicate that the A and B subgenomes show a higher degree of divergence between the three spelt gene pools than the D subgenome, which is also supported by the lower number of D subgenome SNPs obtained from GBS (Table 1). We confirmed that the differences observed in the D subgenome analyses are not a consequence of the lower marker number *per se*. For this, we randomly sub-sampled around 3,500 A and B subgenome-specific SNPs and repeated the PCA (Supporting Information Figure S4). The total variation explained by these sub-samples was similar to the PCAs constructed with all the A and B subgenome markers. Also, we are not aware of any genomic variation in the D subgenome compared to the A and B subgenomes (TE content, gene number) that could explain the reduced diversity as a result of a technical artifact. The ADMIXTURE analysis at K=7 revealed a similar picture for all three subgenomes (Figure 3e). In conclusion, the analysis of individual subgenomes revealed a similar picture as for the whole-genome analysis. The D subgenome, however, is less diverse than the A and B subgenomes.

### Demographic modeling

Considering its geographical proximity, Iberian spelt shows a striking morphological (Figure 1) and genetic (Figure 2) discontinuity with Central European spelt. As a first step towards understanding the origin of Iberian spelt and possible reasons for its peculiar genetic makeup, we compared three evolutionary scenarios with fastsimcoal2, a coalescent simulation tool which allows fitting demographic models to SNP site frequency spectra (Excoffier et al., 2013). In particular, we tested the hypothesis that Iberian spelt was not derived from Central European spelt, but rather brought westwards through a separate migration route.

According to a simple westward migration model (westMig, Figure 4), Iberian and Central European spelt share a common origin and their introduction into Central Europe occurred through a single migration event from the Fertile Crescent. The separation into two gene pools might have been the result of geographic separation, possibly by the Pyrenees. In contrast, in an independent westwards migration model (indepWestMig, Figure 4), Central European and Iberian spelt were brought to Europe independently, possibly through two separate migration corridors north and south of the Alps. Finally, a hybrid model (westMigAdmix, Figure 4) assumes the same stepwise westward migration as westMig, but allows for a recent episode of gene flow from Asian to Iberian spelt. This process could account for the intermediate position of Iberian spelt on the first PCA axis and its morphological similarity to the Asian spelt. In all these models, the ancestral population was either assumed to be the Asian population or an unsampled ghost population, resulting in a total of six models (Figure 4). We fitted the models to two-dimensional SFS estimated from 2,644 non-genic SNPs at least 20 kb apart from each other. Only SNPs in the D subgenome were considered because of the possible polyphyletic origin of the A and B subgenomes.

**Figure 4.**
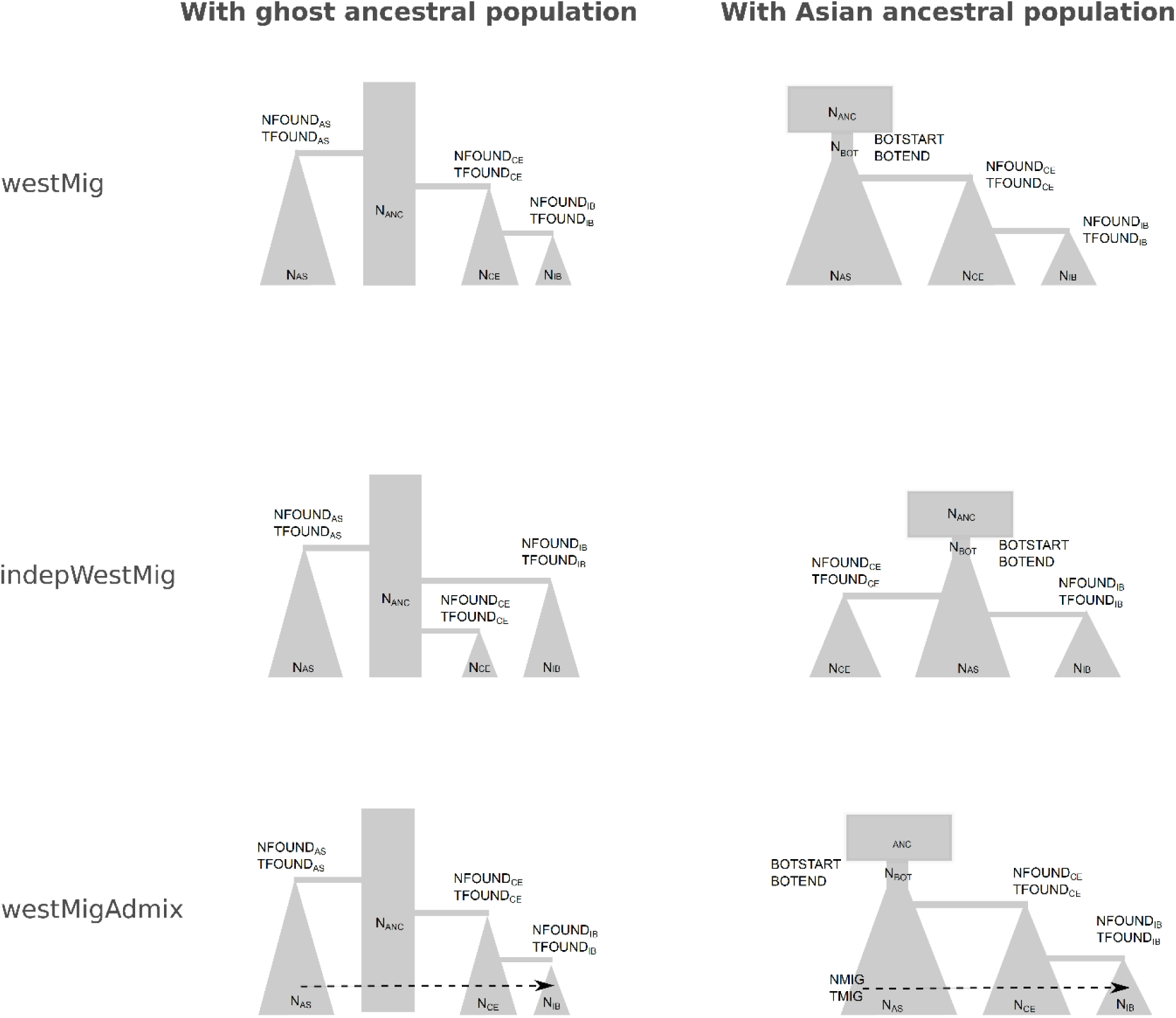
The six demographic models which where compared in this study. Subscripts indicate population names (AS: Asia, CE: Central Europe, IB: Iberia, ANC: ancestral); N indicates population sizes, T indicates times. Detailed parameters are given in Supplementary Information Table S2.

All models agree that Iberian spelt has experienced a stronger bottleneck and has an effective population size which is about half the size of Central European spelt (Supporting Information Table S2). Divergence times of Iberian and Central European spelt from their respective ancestral populations were estimated 1,218 (CI=1,018-1,280) and 1,023 (CI=1,014-1,072) generations ago, respectively. Together with the estimates of effective population size, these split times imply a strong population expansion in the recent past: a 3,665-fold increase compared to the bottlenecked population in the Iberian and a 5,455-fold increase in the Central European population, which is in accordance with recent expansion of agriculture in Europe.

The independent westward migration model consistently attained the best fit to the data: it achieved the highest likelihoods and the lowest AIC values with both ancestral population structures (Figure 4; Supporting Information Table S3), thus supporting the hypothesis that Iberian and Central European spelt have separate migration histories. The second best model was westMig, while the model including recent gene flow from Asian to Iberian spelt (westMigAdmix) performed worst (Supporting Information Table S3). Furthermore, assuming an ancestral ghost population increased the model fit by a large margin, so that the worst ghost population model outperformed the models in which the Asian population was assumed to be ancestral (Supporting Information Table S3). This suggests that European D subgenomes are not straightforward descendants of the Asian spelt accessions sampled in this study, but derived from a spelt population that has not been sampled or that may no longer exist.

In summary, the demographic modeling provides a possible explanation for the striking morphological and genetic differences of Iberian and Central European spelt, as they might be the result of two independent migration events.

## Discussion

Here, we provide a comprehensive population genomic assessment of spelt diversity. Our analyses contribute to the discussions about the definition and origin(s) of spelt.

### What is spelt?

The separation of hexaploid wheat into *aestivum* and *spelta* has mostly been based on morphological spike characteristics. While spelt shows a narrow, elongated spike with brittle rachis and tenacious glumes, bread wheat has a more compact, square spike phenotype with a non-brittle rachis and free-threshing grains. It has been shown that the spike morphology in wheat is controlled by a relatively low number of loci. For example, the *Q* locus located on wheat chromosome arm 5AL represents a major determinant of spike morphology. *Q* codes for an AP2 family transcription factor that pleiotropically influences rachis fragility, glume shape and tenacity, and spike length (Simons et al., 2006). The recessive *q* allele is found in European spelt and confers the typical spelt-like spike. In contrast, the dominant *Q* allele of bread wheat shows higher transcript levels, resulting in more compact spikes with free-threshing seeds. The *Q* allele differs from *q* by one non-synonymous polymorphism in the open reading frame and five conserved polymorphisms in the promoter region (Simons et al., 2006). Interestingly, certain spelt accessions from Asia have been reported to carry the *Q* allele otherwise found in bread wheat (Luo et al., 2000), which led to the hypothesis that *Q* has a regulatory function and that the expression level of *Q*, and hence control of the spike morphology, depends on the genetic background. Another genetic determinant of spike characteristics in cereals is the locus containing the two tightly linked genes *Non-brittle rachis 1* (*btr1*) and *Non-brittle rachis 2* (*btr2*) identified on chromosomes 3 of barley and diploid einkorn (Pourkheirandish et al., 2018; Pourkheirandish et al., 2015). The two genes code for short proteins with no domain homology, and mutations in either of the two genes converts a brittle rachis into a non-brittle one, which is associated with cell wall thickness in the spikelet disarticulation zone. The homoeologous group 2 chromosomes of wheat carry the yet uncloned *tenacious glumes* (*Tg*) genes that control glume tenacity (Dvorak et al., 2012; Faris, Zhang, & Chao, 2014). Furthermore, 15 QTLs for domestication-related spike morphology were characterized in a bi-parental cross between a synthetic hexaploid wheat (durum wheat x *Ae. tauschii*) and the free-threshing bread wheat landrace Chinese Spring (Katkout et al., 2014). Taking all these reports together, it can be concluded that the domestication-related spike characteristics in wheat are controlled by approximately 20 loci.

A taxonomic separation into subspecies can be done based on morphological or genetic differences. Often, the two correlate. In cultivated Asian rice (*Oryza sativa*) for example, several subspecies or varietal groups are distinguished based on morphology and geographical distribution. The two major types are *indica* and *japonica* rice, of which the former has elongated kernels and results in fluffy rice when cooked, while the latter has roundish kernels resulting in sticky rice. Phylogenomic analyses based on whole-genome sequencing showed that the two subspecies also represent two clearly distinct clades (W. S. Wang et al., 2018; Wing, Purugganan, & Zhang, 2018). Similarly, separation of barley landraces into six-rowed and two-rowed, the major morphological determinant of barley spike morphology, was reflected in a clear separation into two gene pools based on exome sequencing (Russell et al., 2016). The situation for spelt though is different. Our whole-genome marker analysis did not result in a separation of a common spelt from the bread wheat gene pool. Asian spelt was indistinguishable from bread wheat based on our molecular analysis, while Central European and Iberian spelt formed two clearly distinct groups. Hence, while a separation into ssp. *spelta* and ssp. *aestivum* might be justified based on morphological characteristics, our molecular analysis does not support this. Instead, we propose to split the hexaploid wheat gene pool into three groups, (*i*) bread wheat/Asian spelt, (*ii*) Central European spelt, and (*iii*) Iberian spelt.

### Where does spelt come from?

Hexaploid wheat evolved in the Fertile Crescent during the Neolithic Revolution. It is thus very likely that Asian spelt represents the most ancient of the three spelt gene pools. The fact that Asian spelt was genetically indistinguishable from bread wheat confirms previous reports of a common, monophyletic origin of Asian spelt and bread wheat. There has been a long-lasting debate whether hulled Asian spelt is ancestral to free-threshing bread wheat or vice versa. Crossing experiments showed that the hybridization of free-threshing *T. turgidum* with *Ae. tauschii* resulted in synthetic hexaploid wheat with spelt-like spike characteristics (McFadden & Sears, 1946). McFadden and Sears therefore concluded that hulled spelt-like hexaploid wheat is ancestral to free-threshing bread wheat. These findings are contradicted by archaeological records suggesting that free-threshing hexaploid wheat predates the hulled form (Nesbitt & Samuel, 1996). However, it can be difficult to distinguish archaeological remains of hulled hexaploid wheat from other hulled species such as tetraploid emmer or diploid *Aegilops*. Given the narrow time window between the first archaeological reports of hulled and free-threshing hexaploid wheat, the question of ancestry will be very difficult to resolve. In contrast to Asian spelt, spelt accessions from Europe markedly differed from Asian spelt/bread wheat. Various reports suggested that Central European spelt emerged from a hybridization of a free-threshing hexaploid wheat with a hulled tetraploid emmer (Blatter et al., 2002, 2004; Dvorak et al., 2012; Liu & Tsunewaki, 1991), a hypothesis that is supported by experimental crosses (Mac Key, 1966). A DNA sequence from a 8,200 year-old charred wheat grain excavated at the Çatalhöyük archaeological settlement in Central Turkey revealed a high similarity to modern Central European spelt (Bilgic, Hakki, Pandey, Khan, & Akkaya, 2016). This might indicate that a Central European-like spelt was already present around 6,200 BC at a site that lies along the migration route from the Fertile Crescent to Europe. It would also indicate that the hybridization that gave rise to Central European spelt occurred somewhere between the Fertile Crescent and Central Turkey, and not in Central Europe as it was often suggested.

If European spelt originated from a hybridization between hexaploid wheat and tetraploid emmer, we would expect that the Asian spelt/bread wheat gene pool differed from Central European spelt in the A and B subgenomes but not in the D subgenome. Surprisingly, our D subgenome analyses revealed a separation of the three individual gene pools, although with a lower genetic diversity than the A and B subgenomes. There are two possible explanations for this: (*i*) European spelt and bread wheat have a polyphyletic origin that involved independent hybridization of a tetraploid wheat with two *Ae. tauschii* accessions, or (*ii*) European spelt and bread wheat have a monophyletic origin but genetic bottlenecks and gene flow resulted in a differentiation of the D subgenomes. Given the high degree of similarity between the D subgenomes of bread wheat and European spelt, which is reflected by the lower number of total SNPs and the shorter distances in the phylogenetic tree, a polyphyletic origin would have involved two very closely related *Ae. tauschii* accessions. It is thus more likely that genetic bottlenecks and gene flow explain the pattern observed in Figure 3. Although spontaneous hybridizations between wheats of different ploidy might occur quite frequently in nature, the generation of viable offspring represent sparse events (Kenan-Eichler et al., 2011). It is thus likely that a very limited number of hybridization events gave rise to Central European spelt. In addition, the effective population size of Central European spelt that found its way into Europe after these hybridizations might have been very small. Gene flow after the separation of Central European spelt and bread wheat could further have expedited a separation of the D subgenomes. Recurrent gene flow from *Ae. tauschii* into hexaploid wheat for example has been demonstrated (J. R. Wang et al., 2013). A very interesting observation is that spelt accessions from the Iberian Peninsula formed a clearly distinct cluster, which raises the question about the origin of Iberian spelt. One possible scenario is that Central European spelt was also introduced into the Iberian Peninsula, where yet another hybridization with a tetraploid emmer resulted in Iberian spelt. Our demographic model comparison, however, suggested that Iberian spelt was introduced into Europe independently from Central European spelt.

Interestingly, a recent population genomics analysis found a genetic difference between ancient farmers from Iberia and Central Europe (Valdiosera et al., 2018), which proposed two independent migrations by Neolithic farmers with slightly different gene pools. The first migration route followed the Danube River into Central Europe and the second route went along the Mediterranean coast into the Iberian Peninsula. It is conceivable that these two ancient human populations also brought along and cultivated cereals of slightly different origin. Interestingly, a difference between Iberian and Eastern European germplasm was also observed for rye, another cereal crop that has its origin in the Fertile Crescent (Parat et al., 2016; Schreiber, Himmelbach, Borner, & Mascher, 2018). Parat et al. (2016) explained this difference with different usage of rye (forage vs. human consumption). Spelt, however, is exclusively used for human consumption both in Iberia and Central Europe. It could thus be that the different end use of rye is a consequence of independently introduced gene pools, and not vice versa. The distinct Iberian spelt gene pool could also be the result of recurrent introductions of spelt at a later point in time, for example during the Umayyad conquest of the Iberian Peninsula between 711 and 788 AD. However, in such a scenario we would have expected to detect admixture between the newly introduced spelt accessions (probably originating in Asia) and the more ancient, Central European-like accessions, which was not supported by our demographic modelling.

## Supporting information

## Acknowledgements

We are grateful to the genebanks of the Leibniz Institute of Plant Genetics and Crop Plant Research (IPK), the Centre for Genetic Resources (CGN), the USDA National Plant Germplasm System, and the Swiss National Genebank for providing seeds of the various accessions. We thank Simon Fluckiger and Helen Zbinden for techincal assistance with DNA extraction and Harold E. Bockelmann from USDA-ARS for valuable discussion on the USDA accessions.

This work was supported by the Swiss Federal Office for Agriculture (BLW) in the framework of NAP-PGREL (national plan of action for the conservation and sustainable utilization of plant genetic resources, project 05-NAP-O34), the University of Zurich and the University Research Priority Program ‘Evolution in Action’, and the King Abdullah University of Science and Technology (KAUST).

## Data accessibility

The raw sequence reads were deposited in the short read archive (SRA) on NCBI under the accession PRJNA498918.

## Author contributions

BK, ACR, TM, and SGK conceived the study. TM designed and processed the spelt collection. MA, CS, and TM performed bioinformatics analyses. MA, BK, CS, ACR and SGK wrote the manuscript and all authors read and approved the final manuscript.

## Reference

Alexander, D. H., Novembre, J., & Lange, K. (2009). Fast model-based estimation of ancestry in unrelated individuals. Genome Research, 19(9), 1655–1664. doi:http://www.genome.org/cgi/doi/10.1101/gr.094052.109

Bertin, P., Gregoire, D., Massart, S., & de Froidmont, D. (2004). High level of genetic diversity among spelt germplasm revealed by microsatellite markers. Genome, 47(6), 1043–1052. doi:https://doi.org/10.1139/g04-065

Bilgic, H., Hakki, E. E., Pandey, A., Khan, M. K., & Akkaya, M. S. (2016). Ancient DNA from 8400 year-old Catalhoyuk wheat: implications for the origin of Neolithic agriculture. PLoS ONE, 11(3), e0151974. doi: https://doi.org/10.1371/journal.pone.0151974

Blatter, R. H. E., Jacomet, S., & Schlumbaum, A. (2002). Spelt-specific alleles in HMW glutenin genes from modern and historical European spelt (Triticum spelta L.). Theoretical and Applied Genetics, 104(2-3), 329–337. doi:https://doi.org/10.1007/s001220100680

Blatter, R. H. E., Jacomet, S., & Schlumbaum, A. (2004). About the origin of European spelt (Triticum spelta L.): allelic differentiation of the HMW Glutenin B1-1 and A1-2 subunit genes. Theoretical and Applied Genetics, 108(2), 360–367. doi:https://doi.org/10.1007/s00122-003-1441-7

Caballero, L., Martin, L. M., & Alvarez, J. B. (2007). Agrobiodiversity of hulled wheats in Asturias (North of Spain). Genetic Resources and Crop Evolution, 54(2), 267–277. doi:https://doi.org/10.1007/s10722-005-4049-8

Cubry, P., Tranchant-Dubreuil, C., Thuillet, A. C., Monat, C., Ndjiondjop, M. N., Labadie, K.,… Vigouroux, Y. (2018). The rise and fall of African rice cultivation revealed by analysis of 246 new genomes. Current Biology, 28(14), 2274–2282. doi:https://doi.org/10.1016/j.cub.2018.05.066

Danecek, P., Auton, A., Abecasis, G., Albers, C. A., Banks, E., DePristo, M. A., … Genomes Project Analysis, G. (2011). The variant call format and VCFtools. Bioinformatics, 27(15), 2156–2158. doi:10.1093/bioinformatics/btr330

Dvorak, J., Deal, K. R., Luo, M. C., You, F. M., von Borstel, K., & Dehghani, H. (2012). The origin of spelt and free-threshing hexaploid wheat. Journal of Heredity, 103(3), 426–441. doi:https://doi.org/10.1093/jhered/esr152

Dwivedi, S. L., Ceccarelli, S., Blair, M. W., Upadhyaya, H. D., Are, A. K., & Ortiz, R. (2016). Landrace germplasm for improving yield and abiotic stress adaptation. Trends in Plant Science, 21(1), 31–42. doi:https://doi.org/10.1016/j.tplants.2015.10.012

Dyck, P. L., & Sykes, E. E. (1994). Genetics of leaf-rust resistance in three spelt wheats. Canadian Journal of Plant Science, 74, 231–233. doi:https://doi.org/10.4141/cjps94-047

Edgar, R. C. (2004). MUSCLE: multiple sequence alignment with high accuracy and high throughput. Nucleic Acids Research, 32(5), 1792–1797. doi:https://doi.org/10.1093/nar/gkh340

Elia, M., Moralejo, M., Rodriguez-Quijano, M., & Molina-Cano, J. L. (2004). Spanish spelt: a separate gene pool within the spelt germplasm. Plant Breeding, 123(3), 297–299. doi:https://doi.org/10.1111/j.1439-0523.2004.00969.x

Eltaher, S., Sallam, A., Belamkar, V., Emara, H. A., Nower, A. A., Salem, K. F. M., … Baenziger, P. S. (2018). Genetic diversity and population structure of F 3:6 Nebraska winter wheat genotypes using genotyping-by-sequencing. Frontiers in Genetics, 9, 76. doi:https://doi.org/10.3389/fgene.2018.00076

Excoffier, L., Dupanloup, I., Huerta-Sanchez, E., Sousa, V. C., & Foll, M. (2013). Robust demographic inference from genomic and SNP data. PLoS Genetics, 9(10), e1003905. doi: https://doi.org/10.1371/journal.pgen.1003905

Faris, J. D., Zhang, Z. C., & Chao, S. M. (2014). Map-based analysis of the tenacious glume gene Tg-B1 of wild emmer and its role in wheat domestication. Gene, 542(2), 198–208. doi:https://doi.org/10.1016/j.gene.2014.03.034

Felsenstein, J. (2005). PHYLIP (Phylogeny Inference Package) version 3.6. Department of Genome Sciences, University of Washington, Seattle.

Glaubitz, J. C., Casstevens, T. M., Lu, F., Harriman, J., Elshire, R. J., Sun, Q., & Buckler, E. S. (2014). TASSEL-GBS: A high capacity genotyping by sequencing analysis pipeline. PLoS ONE, 9(2), e90346. doi: https://doi.org/10.1371/journal.pone.0090346

Gutenkunst, R. N., Hernandez, R. D., Williamson, S. H., & Bustamante, C. D. (2009). Inferring the joint demographic history of multiple populations from multidimensional SNP frequency data. PLoS Genetics, 5(10), e1000695. doi: https://doi.org/10.1371/journal.pgen.1000695

Hunter, J. D. (2007). Matplotlib: a 2D graphics environment. Computing in Science & Engineering, 9(3), 90–95. doi:https://doi.org/10.1109/Mcse.2007.55

International Wheat Genome Sequencing Consortium. (2014). A chromosome-based draft sequence of the hexaploid bread wheat (Triticum aestivum) genome. Science, 345(6194), 1251788. doi:http://dx.doi.org/10.1126/science.1251788

International Wheat Genome Sequencing Consortium. (2018). Shifting the limits in wheat research and breeding through a fully annotated and anchored reference genome sequence. Science, 361, eaar7191. doi:http://dx.doi.org/10.1126/science.aar7191

Katkout, M., Kishii, M., Kawaura, K., Mishina, K., Sakuma, S., Umeda, K., … Ogihara, Y. (2014). QTL analysis of genetic loci affecting domestication-related spike characters in common wheat. Genes & Genetic Systems, 89(3), 121–131. doi:https://doi.org/10.1266/ggs.89.121

Kema, G. H. J. (1992). Resistance in spelt wheat to yellow rust Euphytica, 63(3), 225–231. doi:https://doi.org/10.1007/BF00024548

Kenan-Eichler, M., Leshkowitz, D., Tal, L., Noor, E., Melamed-Bessudo, C., Feldman, M., & Levy, A. A. (2011). Wheat hybridization and polyploidization results in deregulation of small RNAs. Genetics, 188(2), 263–272. doi: https://doi.org/10.1534/genetics.111.128348

Kuckuck, H., & Schiemann, E. (1957). Über das Vorkommen von Speltz und Emmer (Triticum spelta L. und Tr. dicoccum Schubl.) im Iran. Z. Pflanzenzuchtung, 38, 383–396.

Lee, T. H., Guo, H., Wang, X. Y., Kim, C., & Paterson, A. H. (2014). SNPhylo: a pipeline to construct a phylogenetic tree from huge SNP data. Bmc Genomics, 15, 162. doi:https://doi.org/10.1186/1471-2164-15-162

Li, H., & Durbin, R. (2009). Fast and accurate short read alignment with Burrows-Wheeler transform. Bioinformatics, 25(14), 1754–1760. doi:https://doi.org/10.1093/bioinformatics/btp324

Liu, Y. G., & Tsunewaki, K. (1991). Restriction-Fragment-Length-Polymorphism (RFLP) analysis in wheat. II. Linkage maps of the RFLP sites in common wheat. Japanese Journal of Genetics, 66(5), 617–633. doi:https://doi.org/10.1266/jjg.66.617

Luo, M. C., Yang, Z. L., & Dvorak, J. (2000). The Q locus of Iranian and European spelt wheat. Theoretical and Applied Genetics, 100(3-4), 602–606. doi:https://doi.org/10.1007/s001220050079

Ma, J., & Bennetzen, J. L. (2004). Rapid recent growth and divergence of rice nuclear genomes. Proc Natl Acad Sci U S A, 101(34), 12404–12410. doi:https://doi.org/10.1073/pnas.0403715101

Mac Key, J.(1966). Species relationship in Triticum. Paper presented at the Proceedings of the Second International Wheat Genetics Symposium, Lund, Sweden.

Marcussen, T., Sandve, S. R., Heier, L., Spannagl, M., Pfeifer, M., IWGSC, … Olsen, O. A. (2014). Ancient hybridizations among the ancestral genomes of bread wheat. Science, 345(6194), 1250092. doi:http://dx.doi.org/10.1126/science.1250092

McFadden, E. S., & Sears, E. R. (1946). The origin of Triticum spelta and its free-threshing hexaploid relatives. Journal of Heredity, 37(3), 81–89.

McKinney, W. (2010). Data structures for statistical computing in Python. Paper presented at the Proceedings of the 9th Python in Science Conference, Austin, TX.

Meyer, R. S., DuVal, A. E., & Jensen, H. R. (2012). Patterns and processes in crop domestication: an historical review and quantitative analysis of 203 global food crops. New Phytologist, 196(1), 29–48. doi:https://doi.org/10.1111/j.1469-8137.2012.04253.x

Muller, T., Schierscher-Viret, B., Fossati, D., Brabant, C., Schori, A., Keller, B., & Krattinger, S. G. (2018). Unlocking the diversity of genebanks: whole-genome marker analysis of Swiss bread wheat and spelt. Theoretical and Applied Genetics, 131(2), 407–416. doi:https://doi.org/10.1007/s00122-017-3010-5

Nesbitt, M., & Samuel, D. (1996). From staple crop to extinction? The archaeology and history of the hulled wheats. In S. Padulosi, K. Hammer, & J. Heller (Eds.), Hulled wheat - promoting the conservation and use of underutilized and neglected crops 4. Proceedings of the first international workshop on hulled wheats (pp. 40–100). Castelvecchio Pascoli, Tuscany, Italy: International Plant Genetics Resources Institute

Parat, F., Schwertfirm, G., Rudolph, U., Miedaner, T., Korzun, V., Bauer, E., … Tellier, A. (2016). Geography and end use drive the diversification of worldwide winter rye populations. Molecular Ecology, 25(2), 500–514. doi:https://doi.org/10.1111/mec.13495

Pedregosa, F., Varoquaux, G., Gramfort, A., Michel, V., Thirion, B., Grisel, O., … Duchesnay, E. (2011). Scikit-learn: Machine learning in Python. Journal of Machine Learning Research, 12, 2825–2830.

Perez, F., & Granger, B. E. (2007). IPython: a system for interactive scientific computing. Computing in Science & Engineering, 9(3), 21–29. doi:http://dx.doi.org/10.1109/Mcse.2007.53

Poland, J. A., Brown, P. J., Sorrells, M. E., & Jannink, J. L. (2012). Development of high-density genetic maps for barley and wheat using a novel two-enzyme genotyping-by-sequencing approach. PLoS ONE, 7(2), e32253. doi: https://doi.org/10.1371/journal.pone.0032253

Pourkheirandish, M., Dai, F., Sakuma, S., Kanamori, H., Distelfeld, A., Willcox, G., … Komatsuda, T. (2018). On the origin of the non-brittle rachis trait of domesticated einkorn wheat. Frontiers in Plant Science, 8, 2031. doi:https://doi.org/10.3389/fpls.2017.02031

Pourkheirandish, M., Hensel, G., Kilian, B., Senthil, N., Chen, G., Sameri, M., … Komatsuda, T. (2015). Evolution of the grain dispersal system in barley. Cell, 162(3), 527–539. doi:https://doi.org/10.1016/j.cell.2015.07.002

Purcell, S., Neale, B., Todd-Brown, K., Thomas, L., Ferreira, M. A. R., Bender, D., … Sham, P. C. (2007). PLINK: A tool set for whole-genome association and population-based linkage analyses. American Journal of Human Genetics, 81(3), 559–575. doi:https://doi.org/10.1086/519795

Russell, J., Mascher, M., Dawson, I. K., Kyriakidis, S., Calixto, C., Freund, F., … Waugh, R. (2016). Exome sequencing of geographically diverse barley landraces and wild relatives gives insights into environmental adaptation. Nature Genetics, 48(9), 1024–1030. doi:https://doi.org/10.1038/ng.3612

Salamini, F., Ozkan, H., Brandolini, A., Schafer-Pregl, R., & Martin, W. (2002). Genetics and geography of wild cereal domestication in the Near East. Nature Reviews Genetics, 3(6), 429–441. doi:https://doi.org/10.1038/nrg817

Schliep, K. P. (2011). phangorn: phylogenetic analysis in R. Bioinformatics, 27(4), 592–593. doi:https://doi.org/10.1093/bioinformatics/btq706

Schreiber, M., Himmelbach, A., Borner, A., & Mascher, M. (2018). Genetic diversity and relationship between domesticated rye and its wild relatives as revealed through genotyping-by-sequencing. Evolutionary Applications, 1-12. doi:https://doi.org/10.1111/eva.12624

Schreiber, M., Stein, N., & Mascher, M. (2018). Genomic approaches for studying crop evolution. Genome Biology, 19(1), 140. doi:https://doi.org/10.1186/s13059-018-1528-8

Simons, K. J., Fellers, J. P., Trick, H. N., Zhang, Z. C., Tai, Y. S., Gill, B. S., & Faris, J. D. (2006). Molecular characterization of the major wheat domestication gene Q. Genetics, 172(1), 547–555. doi:https://doi.org/10.1534/genetics.105.044727

Sun, Q., Wei, Y., Ni, Z., Xie, C., & Yang, T. (2002). Microsatellite marker for yellow rust resistance gene Yr5 in wheat introgressed from spelt wheat. Plant Breeding, 121(6), 539–541. doi:https://doi.org/10.1046/j.1439-0523.2002.00754.x

Tanksley, S. D., & McCouch, S. R. (1997). Seed banks and molecular maps: unlocking genetic potential from the wild. Science, 277(5329), 1063–1066. doi:https://doi.org/10.1126/science.277.5329.1063

Valdiosera, C., Gunther, T., Vera-Rodriguez, J. C., Urena, I., Iriarte, E., Rodriguez-Varela, R.,… Jakobsson, M. (2018). Four millennia of Iberian biomolecular prehistory illustrate the impact of prehistoric migrations at the far end of Eurasia. Proc Natl Acad Sci U S A, 115(13), 3428–3433. doi:https://doi.org/10.1073/pnas.1717762115

van der Walt, S., Colbert, S. C., & Varoquaux, G. (2011). The NumPy array: a structure for efficient numerical computation. Computing in Science & Engineering, 13(2), 22–30. doi:https://doi.org/10.1109/Mcse.2011.37

Wambugu, P. W., Ndjiondjop, M. N., & Henry, R. J. (2018). Role of genomics in promoting the utilization of plant genetic resources in genebanks. Briefings in Functional Genomics, 17(3), 198–206. doi:https://doi.org/10.1093/bfgp/ely014

Wang, J. R., Luo, M. C., Chen, Z. X., You, F. M., Wei, Y. M., Zheng, Y. L., & Dvorak, J. (2013). Aegilops tauschii single nucleotide polymorphisms shed light on the origins of wheat D-genome genetic diversity and pinpoint the geographic origin of hexaploid wheat. New Phytologist, 198(3), 925–937. doi:https://doi.org/10.1111/nph.12164

Wang, W. S., Mauleon, R., Hu, Z. Q., Chebotarov, D., Tai, S. S., Wu, Z. C., … Leung, H. (2018). Genomic variation in 3,010 diverse accessions of Asian cultivated rice. Nature, 557(7703), 43–49. doi:https://doi.org/10.1038/s41586-018-0063-9

Wing, R. A., Purugganan, M. D., & Zhang, Q. F. (2018). The rice genome revolution: from an ancient grain to Green Super Rice. Nature Reviews Genetics, 19(8), 505–517. doi:https://doi.org/10.1038/s41576-018-0024-z

Wu, G. A., Terol, J., Ibanez, V., Lopez-Garcia, A., Perez-Roman, E., Borreda, C., … Talon, M. (2018). Genomics of the origin and evolution of Citrus. Nature, 554(7692), 311–316. doi:https://doi.org/10.1038/nature25447

Wu, J., Wang, Y. T., Xu, J. B., Korban, S. S., Fei, Z. J., Tao, S. T., … Zhang, S. L. (2018). Diversification and independent domestication of Asian and European pears. Genome Biology, 19, 77. doi:https://doi.org/10.1186/s13059-018-1452-y

Wulff, B. B. H., & Dhugga, K. S. (2018). Wheat-the cereal abandoned by GM. Science, 361(6401), 451–452. doi:https://doi.org/10.1126/science.aat5119

Yoshioka, M., Iehisa, J. C. M., Ohno, R., Kimura, T., Enoki, H., Nishimura, S., … Takumi, S. (2017). Three dominant awnless genes in common wheat: Fine mapping, interaction and contribution to diversity in awn shape and length. PLoS ONE, 12(4), e0176148. doi: https://doi.org/10.1371/journal.pone.0176148

Zheng, X. W., Levine, D., Shen, J., Gogarten, S. M., Laurie, C., & Weir, B. S. (2012). A high-performance computing toolset for relatedness and principal component analysis of SNP data. Bioinformatics, 28(24), 3326–3328. doi:https://doi.org/10.1093/bioinformatics/bts606

