## Supplementary material for "High-throughput genotyping of the spelt gene pool reveals patterns of agricultural history in Europe"

**Online Resource 1: Model parameters**

**# westMig**

//Parameters for the coalescence simulation program : fastsimcoal.exe
4 samples to simulate :
//Population effective sizes (number of genes)
NANC
NSE
NEU
NAS
//Samples sizes and samples age
0 0 0
42 0 0.95
254 0 0.95
24 0 0.95
//Growth rates : negative growth implies population expansion
0
RSE
REU
RAS
//Number of migration matrices : 0 implies no migration between demes
0
//historical event: time, source, sink, migrants, new deme size, new growth rate, migration matrix index
3 historical event
TFOUNDSE 1 2 1 1 REU 0
TFOUNDEU 2 0 1 1 0 0
TFOUNDAS 3 0 1 1 0 0
//Number of independent loci [chromosome]
1 0
//Per chromosome: Number of contiguous linkage Block: a block is a set of contiguous loci
1
//per Block:data type, number of loci, per generation recombination and mutation rates and optional parameters
FREQ 1 0 1.3e-8 OUTEXP

// Priors and rules file
// *********************
[PARAMETERS]
//#isInt? #name #dist.#min #max
//all Ns are in number of haploid individuals
1 NAS unif 1000 1000000 output
1 NEU unif 1000 1000000 output
1 NSE unif 1000 1000000 output
1 NANC unif 1000 10000000 output
1 TFOUNDAS unif 1000 12000 output bounded
1 NFOUNDAS unif 10 100 output
1 TFOUNDSE unif 1000 12000 output bounded
1 NFOUNDSE unif 10 100 output

1 TFOUNDEU unif 1000 12000 output bounded
1 NFOUNDEU unif 10 100 output

[RULES]
TFOUNDSE < TFOUNDEU
TFOUNDEU < TFOUNDAS

[COMPLEX PARAMETERS]
0 SIZERATIOSE = NFOUNDSE/NSE hide
0 LOGSE = log(SIZERATIOSE) hide
0 RSE = LOGSE/TFOUNDSE output
0 SIZERATIOEU = NFOUNDEU/NEU hide
0 LOGEU = log(SIZERATIOEU) hide
0 REU = LOGEU/TFOUNDEU output
0 SIZERATIOAS = NFOUNDAS/NAS hide
0 LOGAS = log(SIZERATIOAS) hide
0 RAS = LOGAS/TFOUNDAS output

**# indepWestMig**

//Parameters for the coalescence simulation program : fastsimcoal.exe
4 samples to simulate :
//Population effective sizes (number of genes)
NANC
NSE
NEU
NAS
//Samples sizes and samples age
0 0 0
42 0 0.95
254 0 0.95
24 0 0.95
//Growth rates : negative growth implies population expansion
0
RSE
REU
RAS
//Number of migration matrices : 0 implies no migration between demes
0
//historical event: time, source, sink, migrants, new deme size, new growth rate, migration matrix index
3 historical event
TFOUNDSE 1 0 1 1 0 0
TFOUNDEU 2 0 1 1 0 0
TFOUNDAS 3 0 1 1 0 0
//Number of independent loci [chromosome]
1 0
//Per chromosome: Number of contiguous linkage Block: a block is a set of contiguous loci
1
//per Block:data type, number of loci, per generation recombination and mutation rates and optional parameters

FREQ 1 0 1.3e-8 OUTEXP

// Priors and rules file
// *********************
[PARAMETERS]
//#isInt? #name   #dist.#min  #max
//all Ns are in number of haploid individuals
1  NAS  unif  1000  1000000  output
1  NEU  unif  1000  1000000  output
1  NSE  unif  1000  1000000  output
1  NANC  unif  1000  10000000  output
1  TFOUNDAS  unif  1000   12000  output  bounded
1  NFOUNDAS  unif  10   100  output
1  TFOUNDSE  unif  1000   12000  output  bounded
1  NFOUNDSE  unif  10   100  output
1  TFOUNDEU  unif  1000   12000  output  bounded
1  NFOUNDEU  unif  10   100  output

[RULES]

[COMPLEX PARAMETERS]
0  SIZERATIOSE = NFOUNDSE/NSE  hide
0  LOGSE = log(SIZERATIOSE)    hide
0  RSE         = LOGSE/TFOUNDSE  output
0  SIZERATIOEU = NFOUNDEU/NEU  hide
0  LOGEU = log(SIZERATIOEU)    hide
0  REU         = LOGEU/TFOUNDEU  output
0  SIZERATIOAS = NFOUNDAS/NAS  hide
0  LOGAS = log(SIZERATIOAS)    hide
0  RAS         = LOGAS/TFOUNDAS  output

**# westMigAdmix**

//Parameters for the coalescence simulation program : fastsimcoal.exe
4 samples to simulate :
//Population effective sizes (number of genes)
NANC
NSE
NEU
NAS
//Samples sizes and samples age
0 0 0
42 0 0.95
254 0 0.95
24 0 0.95
//Growth rates : negative growth implies population expansion
0
RSE
REU
RAS

//Number of migration matrices : 0 implies no migration between demes
0
//historical event: time, source, sink, migrants, new deme size, new growth rate, migration matrix index
4 historical event
TMIG 1 3 MIGPROP 1 RAS 0
TFOUNDSE 1 2 1 1 REU 0
TFOUNDEU 2 0 1 1 0 0
TFOUNDAS 3 0 1 1 0 0
//Number of independent loci [chromosome]
1 0
//Per chromosome: Number of contiguous linkage Block: a block is a set of contiguous loci
1
//per Block:data type, number of loci, per generation recombination and mutation rates and optional parameters
FREQ 1 0 1.3e-8 OUTEXP

// Priors and rules file
// *********************
[PARAMETERS]
//#isInt? #name   #dist.#min  #max
//all Ns are in number of haploid individuals
1  NAS  unif  1000  1000000  output
1  NEU  unif  1000  1000000  output
1  NSE  unif  1000  1000000  output
1  NANC  unif  1000  10000000  output
1  TFOUNDAS  unif  1000   12000  output  bounded
1  NFOUNDAS  unif  10   100  output
1  TFOUNDSE  unif  1000   12000  output  bounded
1  NFOUNDSE  unif  10   100  output
1  TFOUNDEU  unif  1000   12000  output  bounded
1  NFOUNDEU  unif  10   100  output
1  TMIG  unif  10  5000  output  bounded
1  NMIG  unif  1  1000  output  bounded

[RULES]
TMIG < TFOUNDSE
TFOUNDSE < TFOUNDEU
TFOUNDEU < TFOUNDAS

[COMPLEX PARAMETERS]
0  SIZERATIOSE = NFOUNDSE/NSE  hide
0  LOGSE = log(SIZERATIOSE)    hide
0  RSE         = LOGSE/TFOUNDSE  output
0  SIZERATIOEU = NFOUNDEU/NEU  hide
0  LOGEU = log(SIZERATIOEU)    hide
0  REU         = LOGEU/TFOUNDEU  output
0  SIZERATIOAS = NFOUNDAS/NAS  hide
0  LOGAS = log(SIZERATIOAS)    hide
0  RAS         = LOGAS/TFOUNDAS  output
0  MIGRT = RSE*TMIG  hide

0  SOURCESIZE = NSE*(e^MIGRT)  hide
0  MIGPROP = NMIG/SOURCESIZE  hide

### Models with an Asian ancestral population

**# westMig**

//Parameters for the coalescence simulation program : fastsimcoal.exe
3 samples to simulate :
//Population effective sizes (number of genes)
NSE
NEU
NAS
//Samples sizes and samples age
42 0 0.95
254 0 0.95
24 0 0.95
//Growth rates : negative growth implies population expansion
RSE
REU
RAS
//Number of migration matrices : 0 implies no migration between demes
0
//historical event: time, source, sink, migrants, new deme size, new growth rate, migration matrix index
4 historical event
TFOUNDSE 0 1 1 1 REU 0
TFOUNDEU 1 2 1 1 RAS 0
TBOTEND 2 2 0 1 RAS 0
TBOTSTART 2 2 0 RESIZE 0 0
//Number of independent loci [chromosome]
1 0
//Per chromosome: Number of contiguous linkage Block: a block is a set of contiguous loci
1
//per Block:data type, number of loci, per generation recombination and mutation rates and optional parameters
FREQ 1 0 1.3e-8 OUTEXP

// Priors and rules file
// *********************
[PARAMETERS]
//#isInt? #name   #dist.#min  #max
//all Ns are in number of haploid individuals
1  NAS  unif  1000  1000000  output
1  NEU  unif  1000  1000000  output
1  NSE  unif  1000  1000000  output
1  NANC  unif  1000  10000000  output
1  TBOTSTART  unif  1000  12000  output  bounded

1 TBOTEND unif 1000 12000 output bounded
1 NBOT unif 10 100 output
1 TFOUNDSE unif 1000 12000 output bounded
1 NFOUNDSE unif 10 100 output
1 TFOUNDEU unif 1000 12000 output bounded
1 NFOUNDEU unif 10 100 output

[RULES]
TBOTSTART > TBOTEND
TBOTEND > TFOUNDEU
TFOUNDEU > TFOUNDSE

[COMPLEX PARAMETERS]
0 SIZERATIOSE = NFOUNDSE/NSE hide
0 LOGSE = log(SIZERATIOSE) hide
0 RSE = LOGSE/TFOUNDSE output
0 SIZERATIOEU = NFOUNDEU/NEU hide
0 LOGEU = log(SIZERATIOEU) hide
0 REU = LOGEU/TFOUNDEU output
0 SIZERATIOAS = NBOT/NAS hide
0 LOGAS = log(SIZERATIOAS) hide
0 RAS = LOGAS/TBOTEND output
0 RESIZE = NANC/NBOT output

**# indepWestMig**

//Parameters for the coalescence simulation program : fastsimcoal.exe
3 samples to simulate :
//Population effective sizes (number of genes)
NSE
NEU
NAS
//Samples sizes and samples age
42 0 0.95
254 0 0.95
24 0 0.95
//Growth rates : negative growth implies population expansion
RSE
REU
RAS
//Number of migration matrices : 0 implies no migration between demes
0
//historical event: time, source, sink, migrants, new deme size, new growth rate, migration matrix index
4 historical event
TFOUNDSE 0 2 1 1 RAS 0
TFOUNDEU 1 2 1 1 RAS 0
TBOTEND 2 2 0 1 RAS 0
TBOTSTART 2 2 0 RESIZE 0 0
//Number of independent loci [chromosome]
1 0

```
//Per chromosome: Number of contiguous linkage Block: a block is a set of contiguous loci
1
//per Block:data type, number of loci, per generation recombination and mutation rates and optional parameters
FREQ 1 0 1.3e-8 OUTEXP
```

```
// Priors and rules file
// *********************
[PARAMETERS]
//#isInt? #name   #dist.#min  #max
//all Ns are in number of haploid individuals
1  NAS  unif  1000  1000000  output
1  NEU  unif  1000  1000000  output
1  NSE  unif  1000  1000000  output
1  NANC  unif  1000  10000000  output
1  TBOTSTART  unif  1000  12000  output  bounded
1  TBOTEND  unif  1000 12000  output  bounded
1  NBOT  unif  10  100  output
1  TFOUNDSE  unif  1000   12000  output  bounded
1  NFOUNDSE  unif  10   100  output
1  TFOUNDEU  unif  1000   12000  output  bounded
1  NFOUNDEU  unif  10   100  output

[RULES]
TBOTSTART > TBOTEND
TBOTEND > TFOUNDEU
TBOTEND > TFOUNDSE

[COMPLEX PARAMETERS]
0  SIZERATIOSE = NFOUNDSE/NSE  hide
0  LOGSE = log(SIZERATIOSE)    hide
0  RSE        = LOGSE/TFOUNDSE  output
0  SIZERATIOEU = NFOUNDEU/NEU  hide
0  LOGEU = log(SIZERATIOEU)    hide
0  REU        = LOGEU/TFOUNDEU  output
0  SIZERATIOAS = NBOT/NAS  hide
0  LOGAS = log(SIZERATIOAS)    hide
0  RAS        = LOGAS/TBOTEND  output
0  RESIZE = NANC/NBOT  output
```

**# westMigAdmix**

```
//Parameters for the coalescence simulation program : fastsimcoal.exe
3 samples to simulate :
//Population effective sizes (number of genes)
NSE
NEU
NAS
//Samples sizes and samples age
42 0 0.95
```

254 0 0.95
24 0 0.95
//Growth rates : negative growth implies population expansion
RSE
REU
RAS
//Number of migration matrices : 0 implies no migration between demes
0
//historical event: time, source, sink, migrants, new deme size, new growth rate, migration matrix index
5 historical event
TMIG 0 2 MIGPROP 1 RAS 0
TFOUNDSE 0 1 1 1 REU 0
TFOUNDEU 1 2 1 1 RAS 0
TBOTEND 2 2 0 1 RAS 0
TBOTSTART 2 2 0 RESIZE 0 0
//Number of independent loci [chromosome]
1 0
//Per chromosome: Number of contiguous linkage Block: a block is a set of contiguous loci
1
//per Block:data type, number of loci, per generation recombination and mutation rates and optional parameters
FREQ 1 0 1.3e-8 OUTEXP

// Priors and rules file
// ********************
[PARAMETERS]
//#isInt? #name #dist.#min #max
//all Ns are in number of haploid individuals
1 NAS unif 1000 1000000 output
1 NEU unif 1000 1000000 output
1 NSE unif 1000 1000000 output
1 NANC unif 1000 10000000 output
1 TBOTSTART unif 1000 12000 output bounded
1 TBOTEND unif 1000 12000 output bounded
1 NBOT unif 10 100 output
1 TFOUNDSE unif 1000 12000 output bounded
1 NFOUNDSE unif 10 100 output
1 TFOUNDEU unif 1000 12000 output bounded
1 NFOUNDEU unif 10 100 output
1 TMIG unif 10 5000 output bounded
1 NMIG unif 1 1000 output bounded

[RULES]
TMIG < TFOUNDSE
TFOUNDSE < TFOUNDEU
TFOUNDEU < TBOTEND
TBOTEND < TBOTSTART

[COMPLEX PARAMETERS]
0 SIZERATIOSE = NFOUNDSE/NSE hide

```
0  LOGSE = log(SIZERATIOSE)    hide
0  RSE        = LOGSE/TFOUNDSE  output
0  SIZERATIOEU = NFOUNDEU/NEU  hide
0  LOGEU = log(SIZERATIOEU)    hide
0  REU        = LOGEU/TFOUNDEU  output
0  SIZERATIOAS = NBOT/NAS  hide
0  LOGAS = log(SIZERATIOAS)    hide
0  RAS        = LOGAS/TBOTEND  output
0  RESIZE = NANC/NBOT  output
0  MIGRT = RSE*TMIG  hide
0  SOURCESIZE = NSE*(e^MIGRT)  hide
0  MIGPROP = NMIG/SOURCESIZE  hide
```
