## Supplementary material for "High-throughput genotyping of the spelt gene pool reveals patterns of agricultural history in Europe"

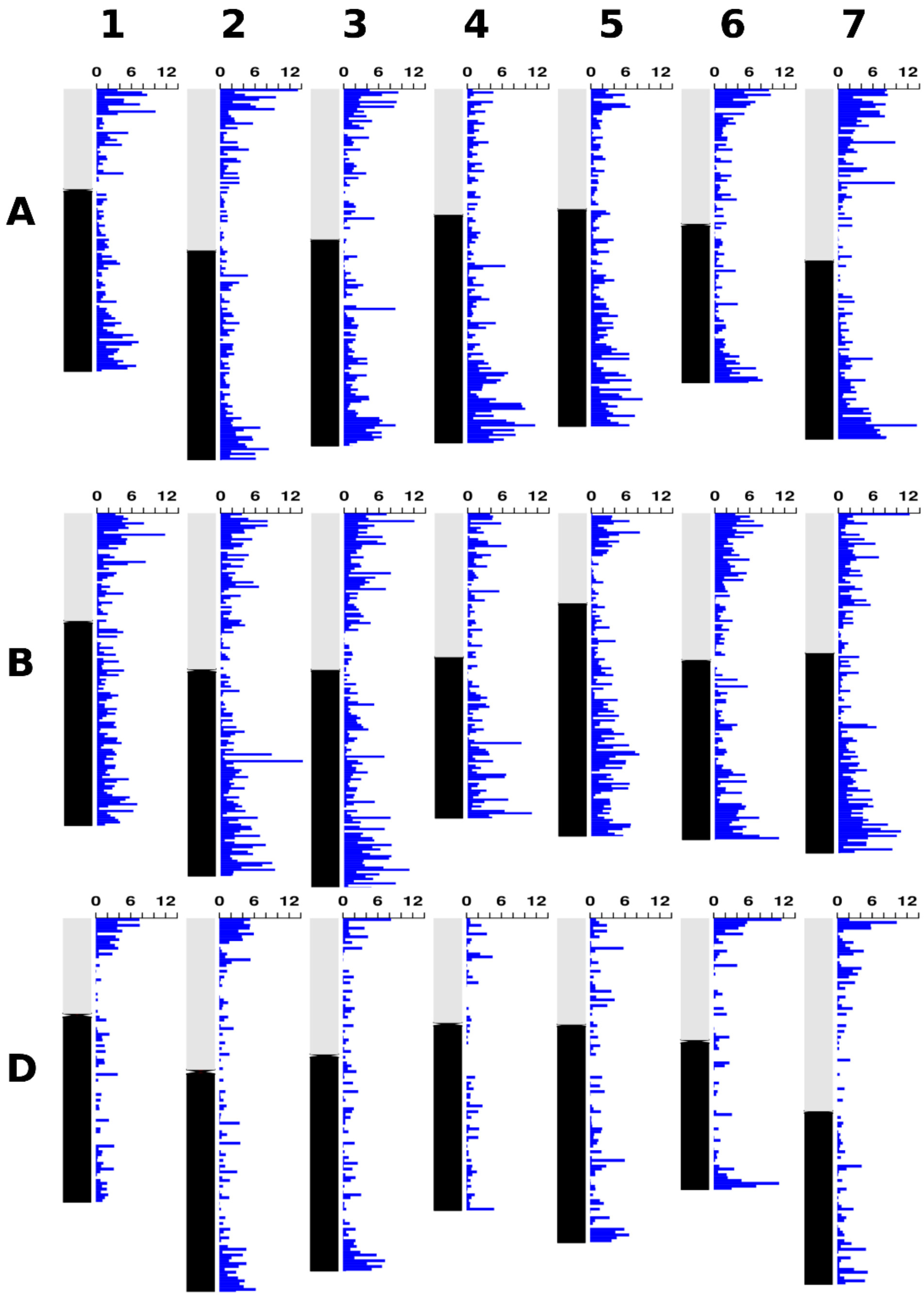


**Figure S1.** Distribution of SNP markers across the different wheat chromosomes. Blue horizontal bars indicate the number of SNP markers in a sliding window of 5 Mb. Short chromosome arms are indicated as vertical grey bars and long chromosome arms are indicated as vertical black bars.

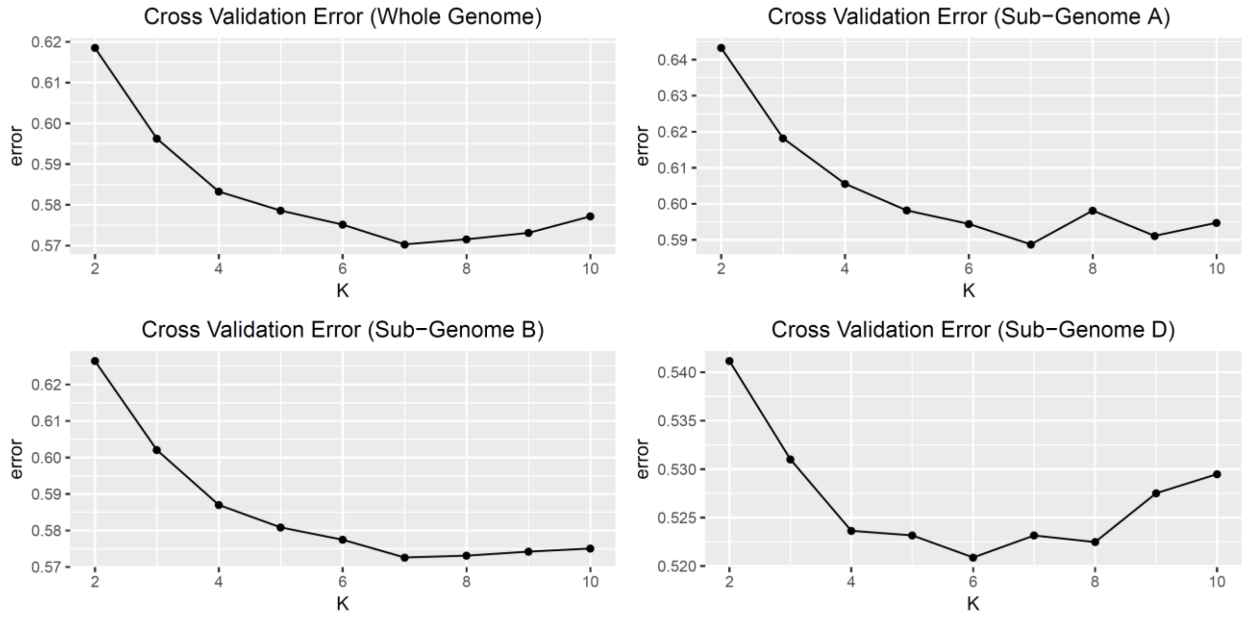


**Figure S2.** Cross validation error plots for the ADMIXTURE analyses of the whole genome and subgenomes A, B and D, respectively.

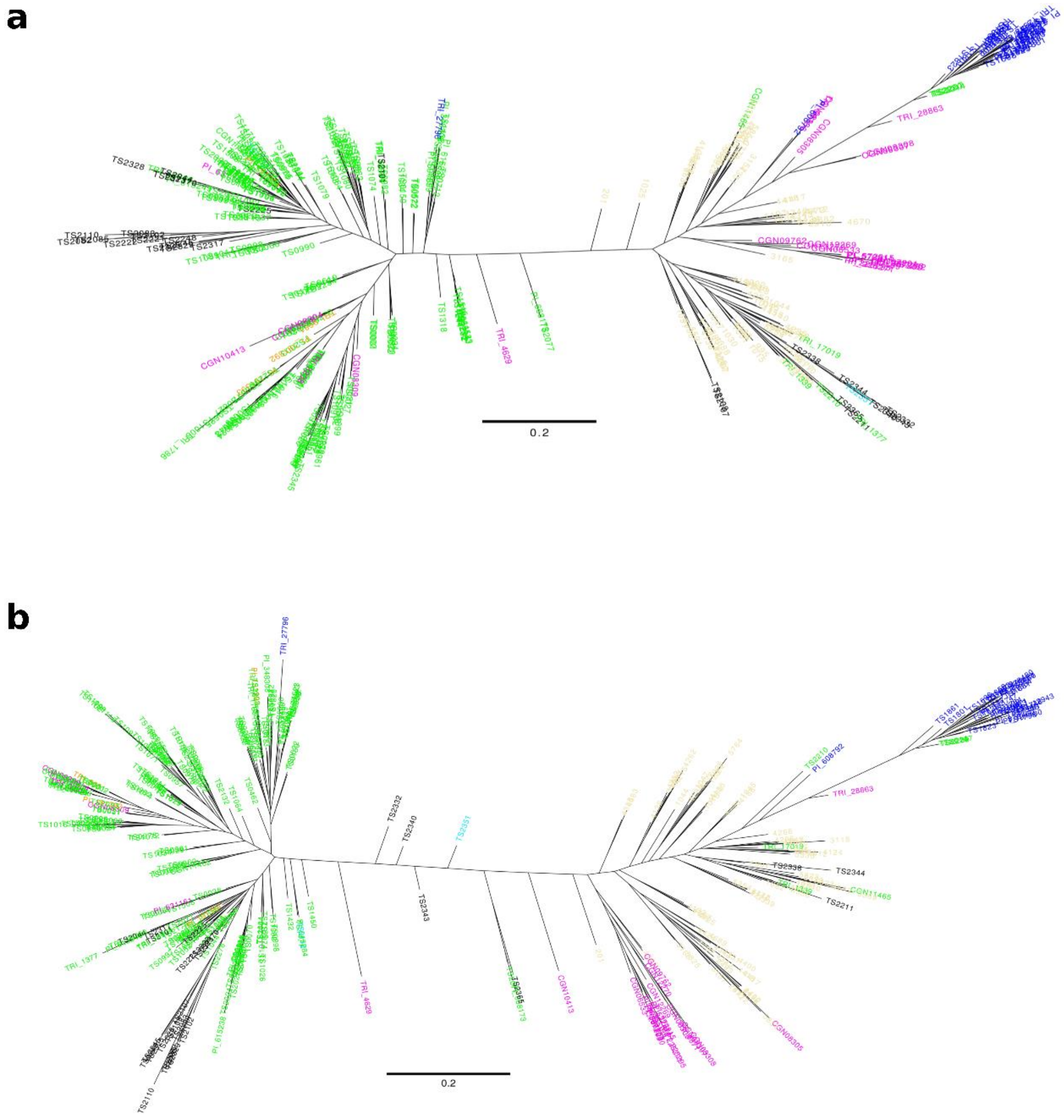


**Figure S3.** Maximum-likelihood trees constructed with SNPhylo for subgenomes A **(a)** and B **(b)**. The colors used to label the accessions are identical to Fig. 2 and Fig. 3. The branch sizes are indicated below the trees.

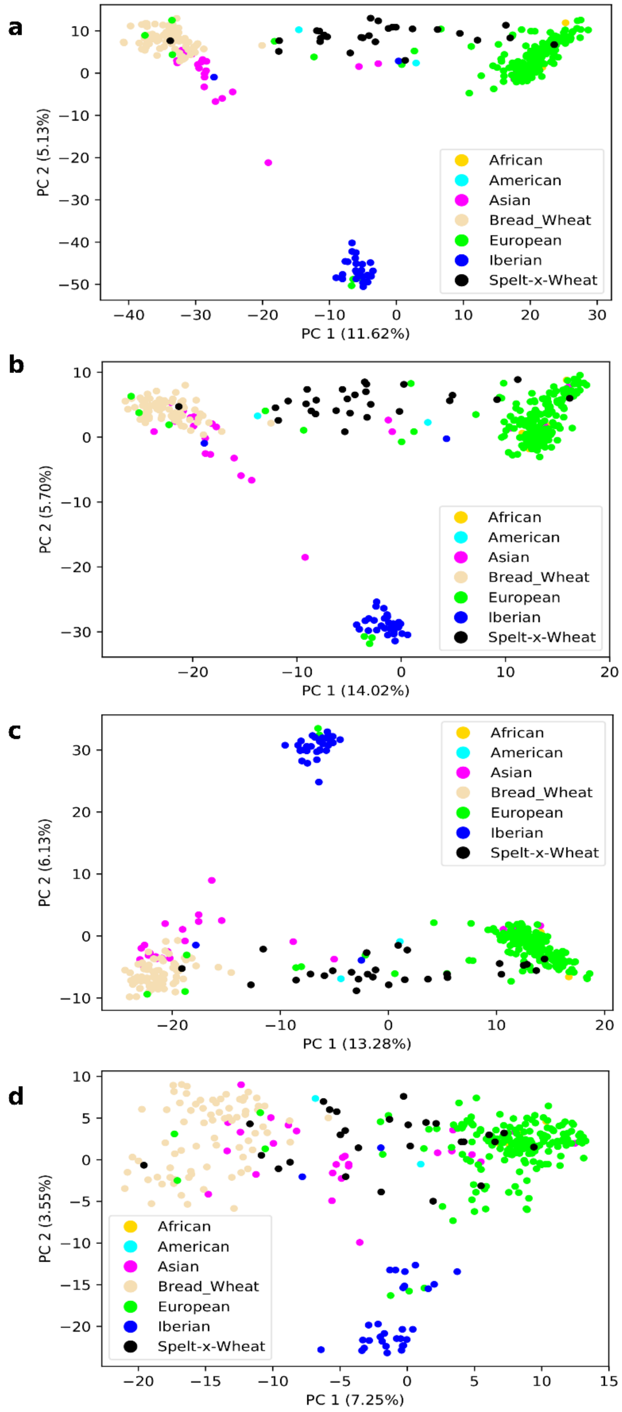


**Figure S4.** Principal component analyses (PCA) with marker sub-sets. PCAs were constructed using 3,500 randomly extracted SNPs of the whole genome **(a)** A subgenome **(b)**, B subgenome **(c)** and D subgenome **(d)**, respectively.

**Table S2.** Parameter estimates for demographic modelling.

**A) Estimates for the best run of all six models**

| | **indepWestMig** | | **westMig** | | **westMigAdmix** | |
|---|---|---|---|---|---|---|
| | Ghost | Asian ancestral | Ghost | Asian ancestral | Ghost | Asian ancestral |
| $N_{AS}$ | 1,036,691 | 1,745,686 | 6,828,368 | 936,104 | 7,199,056 | 1,001,769 |
| $N_{CE}$ | 5,274,951 | 908,584 | 2,976,393 | 2,295,759 | 2,702,867 | 1,503,286 |
| $N_{IB}$ | 2,217,462 | 583,331 | 1,216,222 | 1,004,393 | 1,431,178 | 774,210 |
| $N_{ANC}$ | 29,296,457 | 74,944,876 | 31,506,118 | 65,144,251 | 33,369,192 | 66,837,240 |
| TBOTSTART | NA | 8,121 | NA | 6,684 | NA | 8,444 |
| TBOTEND | NA | 8,000 | NA | 6,610 | NA | 8,357 |
| NBOT | NA | 672 | NA | 591 | NA | 598 |
| $TFOUND_{AS}$ | 1,014 | NA | 5,588 | NA | 8,486 | NA |
| $NFOUND_{AS}$ | 702 | NA | 627 | NA | 547 | NA |
| $TFOUND_{IB}$ | 1,218 | 1,202 | 1,038 | 1,322 | 1,268 | 1,264 |
| $NFOUND_{IB}$ | 605 | 320 | 395 | 318 | 443 | 265 |
| $TFOUND_{CE}$ | 1,023 | 1,039 | 1,074 | 1,395 | 1,293 | 1,299 |
| $NFOUND_{CE}$ | 967 | 458 | 662 | 409 | 696 | 362 |
| $R_{IB}$ | -0.0067 | -0.0062 | -0.0077 | -0.0061 | -0.0064 | -0.0063 |
| $R_{CE}$ | -0.0084 | -0.0073 | -0.0078 | -0.0062 | -0.0064 | -0.0064 |
| $R_{AS}$ | -0.0072 | -0.0010 | -0.0017 | -0.0011 | -0.0011 | -0.0009 |
| TMIG | NA | NA | NA | NA | 958 | 1,103 |
| NMIG | NA | NA | NA | NA | 453 | 700 |

**B) Parameter estimates and confidence intervals for the best model (indepWestMig_ghost)**

| Parameters | Point estimate | Q5% | Q95% |
|---|---|---|---|
| $N_{AS}$ | 1,036,691 | 871,467 | 4,762,619 |
| $N_{CE}$ | 5,274,951 | 795,293 | 4,640,651 |
| $N_{IB}$ | 2,217,462 | 1,676,913 | 3,720,851 |
| $N_{ANC}$ | 29,296,457 | 19,541,580 | 23,011,428 |
| $TFOUND_{AS}$ | 1,014 | 1,011 | 1,069 |
| $NFOUND_{AS}$ | 702 | 439 | 777 |
| $TFOUND_{IB}$ | 1,218 | 1,018 | 1,280 |
| $NFOUND_{IB}$ | 605 | 438 | 617 |
| $TFOUND_{CE}$ | 1,023 | 1,014 | 1,072 |
| $NFOUND_{CE}$ | 967 | 328 | 816 |
| $R_{IB}$ | -0.0067 | -0.0084 | -0.0065 |
| $R_{CE}$ | -0.0084 | -0.0085 | -0.0074 |
| $R_{AS}$ | -0.0072 | -0.0086 | -0.0071 |

In the parameter names, subscripts indicate population names (AS: Asia, CE: Central Europe, IB: Iberia, ANC: ancestral); parameters beginning with N indicate population sizes, those beginning with T times, and those with R population growth rates (which are negative because in the coalescence framework inference goes backwards in time from the present to the past).

**Table S3.** Demographic model comparisons.

| **Model name** | **logL[a]** | **Nr. parameters[b]** | **AIC[c]** |
|---|---|---|---|
| indepWestMig_ghost | -7,758.08 | 10 | **15,536.17** |
| westMig_ghost | -7,983.47 | 10 | **15,986.94** |
| westMigAdmix_ghost | -8,081.42 | 12 | **16,186.84** |
| indepWestMig_AsianAncestral | -9,045.23 | 11 | **18,112.46** |
| westMig_AsianAncestral | -9,176.74 | 11 | **18,375.49** |
| westMixAdmix_AsianAncestral | -9,349.66 | 13 | **18,725.32** |

[a] Log-likelihood of the model. [b] Number of parameters in the model. [c]Akaike information criterion, calculated as 2*Nr. parameters - 2*logL.
